# scPred: Cell type prediction at single-cell resolution

**DOI:** 10.1101/369538

**Authors:** José Alquicira-Hernández, Anuja Sathe, Hanlee P Ji, Quan Nguyen, Joseph E Powell

## Abstract

Single-cell RNA sequencing has enabled the characterization of highly specific cell types in many human tissues, as well as both primary and stem cell-derived cell lines. An important facet of these studies is the ability to identify the transcriptional signatures that define a cell type or state. In theory, this information can be used to classify an unknown cell based on its transcriptional profile; and clearly, the ability to accurately predict a cell type and any pathologic-related state will play a critical role in the early diagnosis of disease and decisions around the personalized treatment for patients. Here we present a new generalizable method (*scPred*) for prediction of cell type(s), using a combination of unbiased feature selection from a reduced-dimension space, and machine-learning classification. *scPred* solves several problems associated with the identification of individual gene feature selection, and is able to capture subtle effects of many genes, increasing the overall variance explained by the model, and correspondingly improving the prediction accuracy. We validate the performance of `scPred` by performing experiments to classify tumor versus non-tumor epithelial cells in gastric cancer, then using independent molecular techniques (cyclic immunohistochemistry) to confirm our prediction, achieving an accuracy of classifying the disease state of individual cells of 99%. Moreover, we apply *scPred* to scRNA-seq data from pancreatic tissue, colorectal tumor biopsies, and circulating dendritic cells, and show that *scPred* is able to classify cell subtypes with an accuracy of 96.1-99.2%. Collectively, our results demonstrate the utility of *scPred* as a single cell prediction method that can be used for a wide variety of applications. The generalized method is implemented in software available here: https://github.com/IMB-Computational-Genomics-Lab/scPred/

## Introduction

Individual cells are the basic building blocks of organisms, and while a human consists of an estimated three trillion cells, each one of them is unique at a transcriptional level. Performing bulk or whole-tissue RNA sequencing, which combines the contents of millions of cells, masks most of the differences between cells as the resulting data comprises of the averaged signal from all cells. Single cell RNA-sequencing (scRNA-seq) has emerged as a revolutionary technique, which can be used to identify the unique transcriptomic profile of each cell. Using this information we are now able to address questions that previously could not be answered, including the identification of new cell types [1–4], resolving the cellular dynamics of developmental processes [5–8]), and identify gene regulatory mechanisms that vary between cell subtypes [9].

Cell type identification and discovery of subtypes has emerged as one of the most important early applications of scRNA-seq [10]. Prior to the arrival of scRNA-seq, the traditional methods to classify cells were based on microscopy, histology, and pathological criteria [11]. In the field of immunology, cell surface markers have been widely used to distinguish cell subtypes [12], for a wide range of purposes. While this approach is desirable in practical terms for cell isolation -*e.g.* via fluorescence-activated cell sorting (FACS)-these markers may not reflect the overall heterogeneity at a transcriptomic and phenotypic level from mixed cell populations [13, 14]. Using scRNA-seq data unsupervised or supervised clustering approaches have been used to determine groups of cells based on similar transcriptional signatures relative the remaining sample [15–17], often followed by functional validation to classify specific cell subtypes. Similarly, differential expression between groups of cells can identify lists cell-type specific markers, and functional enrichment analyses used to help define cell subtype function [18]. These approaches are being used to classify cells subtypes in a given sample, but in an independent sample simple methods to accurately classify each cell are needed.

To be able to predict the classification of a single cell based upon its transcriptome read-out, first a prediction model needs to be built where the effects of given features are estimated. It is clear that both the selection of features, and estimation of their effects plays a critical role in the overall prediction performance. Unlike prediction methods that use data derived from bulk RNA-seq data where gene expression averages are used as features, phenotype prediction at single cell level faces new challenges. Firstly, cell-to-cell differences must be taken into account to define and predict cell types. Using only a subset of genes (e.g. differentially expressed genes) will likely exclude discriminant sources of variation across cells. An additional limitation is the inconsistency seen between statistical methods used to identify differentially expressed genes [19]. Finally, if the number of observations that define a specific subtype of cells is high, then classification algorithms can be computationally expensive, or suffer from overfitting.

There are numerous applications for which prediction of a cell state or type from its scRNA-seq data can play an important role. An obvious example is in the burgeoning use of single cell data in characterizing disease states and underlying biology at single cell resolution [20] [12]. The granular nature of single cell characterization has enormous implications for accurate prediction of specific cell subtypes and pathologic-related states. We anticipate that such prediction strategies will play an important role in early diagnosis of diseases or informing personalized treatment. Similarly, efforts arising from the Human Cell Atlas [10] are set to create a comprehensive reference atlases of most cell subtypes in the human body, meaning cells from new samples can be mapped against this atlas. Here, we introduce `scPred`, a method that takes advantage of dimensionality reduction and orthogonalization of gene expression values to accurately predict specific cell types or states of single cells from their transcriptional data (Figure 1). *scPred* can be applied to any situation where cells can be labeled into discrete categories, including cell subtypes or defined cell states.

**Figure 1:**
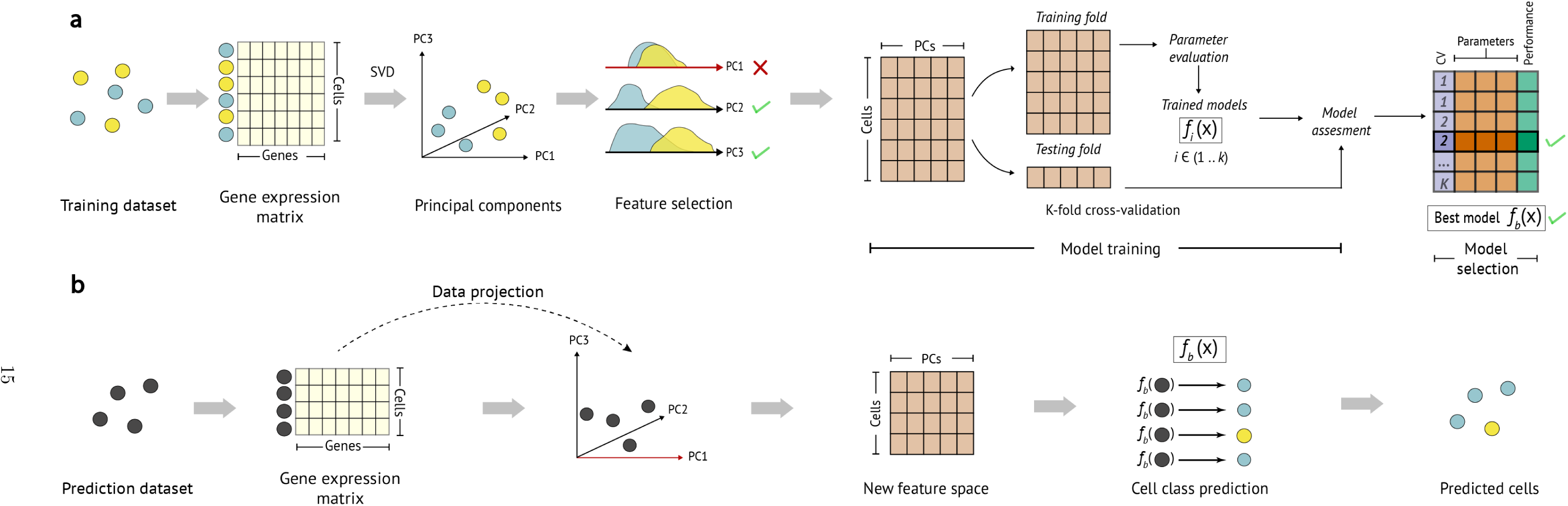
Summary of the `scPred` method. A) Training step. A gene expression matrix is eigendecomposed via singular value decomposition (SVD) to obtain orthonormal linear combinations of the gene expression values. Only PCs explaining greater than 0.01 percent of the variance of the dataset are considered for the feature selection and model training steps. Informative PCs are selected using a two-tailed Wilcoxon signed rank test for each cell class distribution (Methods). The cells-PCs matrix is randomly split into *k* groups and the first *k* group is considered as a testing dataset for cross-validation. The remaining *K-1* groups (shown as a single *training fold*) are used to train a machine learning classification model (a support vector machine). The model parameters are tuned and each k group is used as a testing dataset to evaluate the prediction performance of a *f_i_*(*x*) model trained with the remaining *K-1* groups. The best model in terms of prediction performance is selected. B) Prediction step. The gene expression values of the cells from an independent test or validation dataset are projected onto the principal component basis from the training model, and the informative PCs are used to predict the class probabilities of each cell using the trained prediction model(s) *f_b_*(*x*).

## Results

*scPred* is a generalized method for classifying a single cell based on its transcriptional data. The method uses a combination of decomposing the variance structure of a gene expression matrix to identify limited informative features, and a machine learning approach to estimate the effect of these features on classifying cells (Figure 1). In doing so, it is able to incorporate a large number of small differences in the mean and variance of gene expression between different cell types in the prediction model. This removes the need to perform gene-specific analyses such as differential expression to identify informative features. *scPred* has two main steps. Firstly, a prediction model is built using a training cohort of single cell data, where the identity of the cells is already known. Secondly, the application of the prediction model to single cell data obtained from independent sample, with each cell then assigned a conditional class probability *Pr*(*y* = 1|*f*) of belonging to a given cell subtype or state. Here we present results of the application of *scPred* under three distinct scenarios.

### *scPred* can accurately predict tumor epithelial cells from gastric cancer

To evaluate the performance of scPred we initially sought an orthogonal molecular technique that is able to confirm or reject the probabalistic predictions made by *scPred.* We obtained surgical biopsies from stage IIA intestinal gastric adenocarcinoma along with matched-normal epithelium from two patients, and generated scRNA-seq data using the Chromium platform (10X Genomics). From the a total of 3,921 cells sequenced, we identified 1,905 epithelial cells based on the expression of EpCAM. We split the cells into two sets, a training set comprising of 953 randomly selected cells, leaving 952 cells for prediction. A *scPred* model was developed using the training data, and applied to each of the 952 cells in the prediction set to cell to classify them as either a tumor or non-tumor cell. The malignancy status of predicted cells was confired based on the observed loss of MLH1 and PMS2 protein expression with immunohistochemistry; these DNA mismatch repair proteins form the MutLα heterodimer [21]. Given this characterization, this gastric tumor was classified as being of the molecular subtype characterized by microsatellite instability (MSI) that is a hypermutable state related to loss of DNA mismatch repair proteins such as MLH1. To account for the heterogeneity in the samples, we repeated this step ten times with random splits. Overall, we obtained a mean sensitivity of 97.9% and a specificity of 100% across the ten bootstrap replicates (Figure 2).

**Figure 2:**
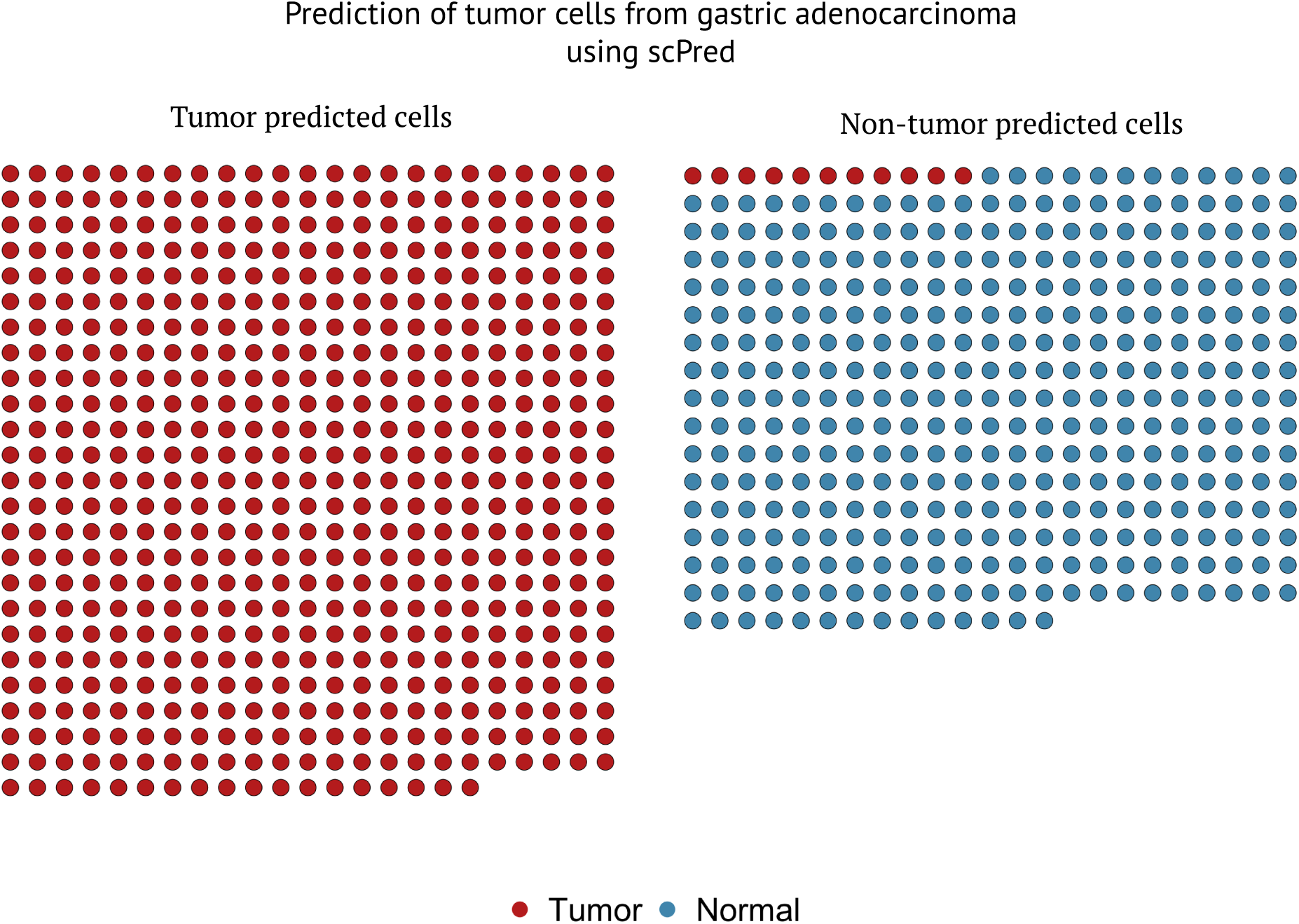
Prediction results of tumor gastric cancer cells. Cells are grouped based on their predicted status by `scPred`. Disease status of each cell (confirmed by the loss of expression of MLH1 and PMS2) is represented by color. See legend.

### *scPred* can accurately predict cell subtype using scRNA-seq data generated across different platforms

Given the rapid development of single cell sequencing assays and technologies, we anticipate that a prediction model for a given cell subtype(s) will often be built with data generated from an alternative platform to that used for independent test samples. To assess the robustness of *scPred* we sought to evaluate the performance using training data generated from multiple platforms, and testing the prediction accuracy for independent cells sequenced on another platform. We chose to develop a prediction model using *scPred* to classify subtypes of islets of Langerhans cells from scRNA-seq data generated from pancreas tissue due to their limited abundance (4.5% in a pancreatic tissue sample) [22], and thus will represent a class of cells that is expected to be more difficult to predict based on their low relative existence compared to other cells.

Islets of Langerhans are composed mainly of four distinct cell types, namely *α* (alpha), *β* (beta), *δ* (delta) and *γ* (gamma) cells, that are responsible of producing glucagon, insulin, somatostatin, and pancreatic-polypeptides respectively [23]. We generated a training reference cohort of scRNA-seq data from a total of 4,292 cells from three independent studies undertaken by Muraro *et al* [24], Segerstolpe *et al* [3], and Xin *et al* [25] that had sequenced cells using CEL-seq2 [26], Smart-Seq2 [27], and SMARTer [28] respectively. Details of the training cohort data are given in Supporting Material Table 1. Importantly, using the Seurat canonical correlation alignment method [29], we are able to demonstrate that between platform and between sample batch effects can be removed for the training cohort (Figure 3). The best fit models from *scPred* for *α*, *β*, *δ* and *γ* cells used between 14-18 PCs, which represents a small feature space for prediction in an independent data, and correspondingly will reduce the computational requirements of *scPred* in the testing phase.

**Table 1:** Prediction of pancreatic cells. The training panel correspond to the *Muraro*, *Segerstolpe* and *Xin* datasets used as a reference to train the prediction models for each cell type from the islets of Langerhans. As part of the training, no other cell types were considered. The test information corresponds to the Baron [30] dataset used to measure the performance of the trained models in a independent dataset. The Baron dataset contains epsilon, acinar, stellate, ductal, endothelial, Schwann and T cells referred as “Other” in this table. The accuracy is defined as the fraction of cells correctly assigned for each cell type of interest. The accuracy for the remaining cells corresponds to the fraction of cells from the test dataset that are correctly unassigned to any of the classes of interest (negative controls) as a consensus across all four prediction models.

| Cell type | Training |  |  | Test |  |
| --- | --- | --- | --- | --- | --- |
|  | # cells | # PCs | # Support vectors | # Cells | Accuracy |
| $\alpha$ Alpha | 2584 | 18 | 362 | 2302 | 98.3 |
| $\beta$ Beta | 1190 | 17 | 343 | 2454 | 96.1 |
| $\delta$ Delta | 356 | 14 | 283 | 596 | 97.1 |
| $\gamma$ Gamma | 383 | 15 | 215 | 254 | 99.2 |
| Other | 0 | NA | NA | 2326 | 94.9 |

**Figure 3:**
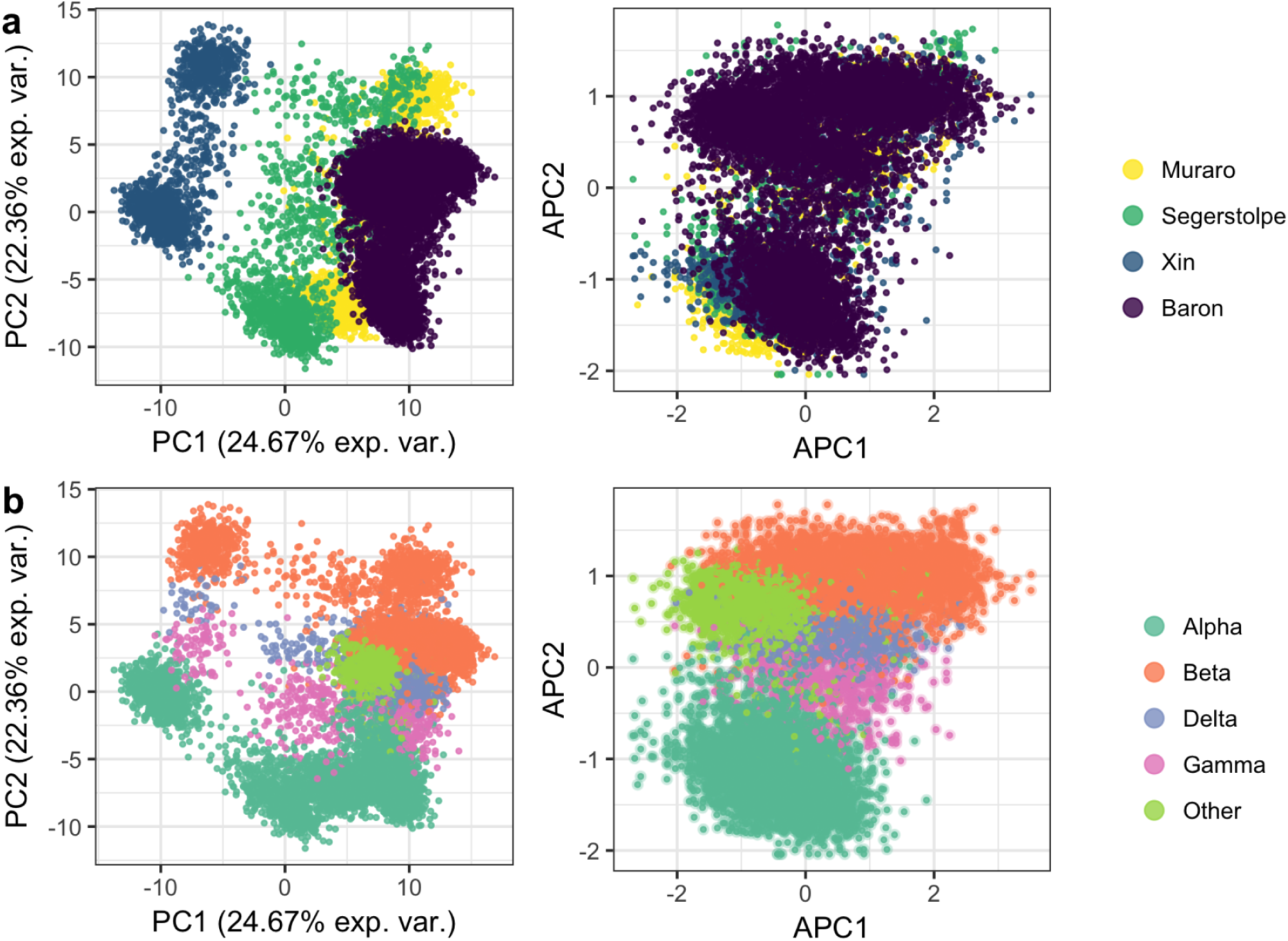
Principal component alignment of pancreatic cells. (a) Training (Muraro, Segerstolpe, and Xin) datasets [3, 24, 25] were used to generate the training eigenspace. The test dataset (Baron [30]) was projected and all datasets were aligned using Seurat. No batch effect is observed after the alignment. (b) *α*, *β*, *δ* and *γ* cells are included in the training datasets. The prediction dataset contains also 2,326 “other” cell types such as epsilon, acinar, stellate, ductal, endothelial, Schwann and T cells (bright green cells). After the dataset alignment, cells cluster by cell type. The *X*-axis shows variance explained (exp.var.), principal components (PC), and aligned principal components (APC).

**Figure 4:**
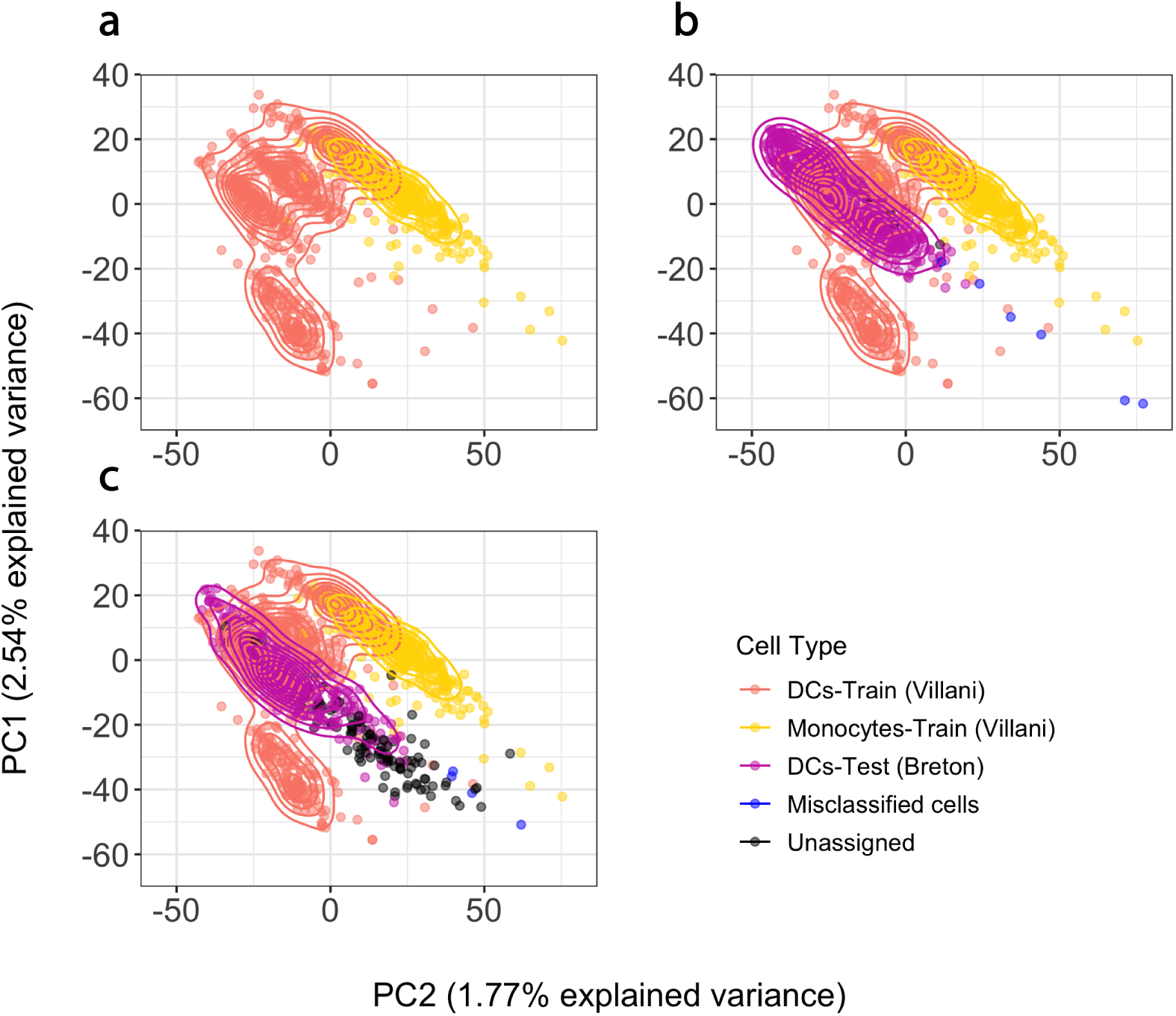
Prediction of human dendritic cells. (a) The training dataset (*Villani et al*) of dendritic cells and monocytes was eigendecomposed (orange and yellow points and density lines). (b) Dendritic cells from the test dataset (*Breton et al*) were projected onto the training eigenspace (purple points). `scPred` predicted 98% of dendritic cells derived from peripheral blood correctly and 82% from umbilical cord (*Breton et al*). Blue points correspond to cells that were misclassified and black points to unassigned cells.

Using the prediction classifier model trained from the reference cohort data, we naively predicted the cell type of each of 7,932 cells [30], collected from four healthy individuals, using their scRNA-seq data generated using inDrop [31]. The testing data includes a heterogeneous mix cells islets of Langerhans cells, meaning non-*α*, *β*, *δ* and *γ* cells, such as the epsilon, endothelial or T cells provide a negative control. We classified a cell as a specific cell subtype based on a class probability (*Pr*(*y* = 1|*f*)) greater than 0.9. The overall accuracy of the predictions were evaluated based on the known cell identities determined based on the expression of classic markers (*GCG, INS, SST,* and *PPY*). For islets of Langerhans cells, the prediction model built by *scPred* using the scRNA-seq data from the reference cohort, was able to predict cell type with an average accuracy of 97.68% (Table 1 and Figure 3) and accurately labeling 94.9% heterogeneous populations of other cells. For example, of the 2,302 *α* cells in the test cohort our *scPred* model classified 2,264 cells correctly. Of the 38 miss-classified cells, 33 were unassigned to another target cell type, which also demonstrates a high specificity of the model. We observed the same pattern for all cell types tested (Supporting Material Table 2). To further support this conclusion, the mean *Pr*(*y* = 1|*f*) for cells classified as *α*, *β*, *δ* and *γ* was 0.994-0.997, while cells classified as other (i.e. epsilon, endothelial or T cells) had a mean *Pr*(*y* = 1|*f*) of 0.307 (Supporting Material Figure 1). These results show that the features selected from the decomposed training data are able to define hyperplanes that are able to separate individual cells by cell type, based upon linear combinations of scRNA-seq data fitted to a *scPred* models.

### Accurate prediction of human dendritic cells from data generated across laboratories

We next sought to evaluate the performance of *scPred* when the training and testing cells sequenced using the same protocol, but in different laboratories. For developing single cell-based diagnostic tests this is an important consideration, as in the majority of cases a predictive model will be developed using sequence data generated from different laboratories to those conducting testing. Between site effects could bias the predictive performance of a test if the between site batch effects are confounded with the model classification features. While between site variance for bulk-RNA-sequencing is small [32], it has not yet been fully evaluated for scRNA-Seq.

We chose to evaluate the performance of *scPred* by building a prediction model to identify dendritic cells from peripheral blood samples [1]. Dendritic cells are antigen-presenting cells, and their main function is to process antigen material and present it on the cell surface to T-cells, acting as messengers between the innate and the adaptive immune systems. Using the cell type classification based on scRNA-seq and flow validation given in Villani *et al.* [1], we built a *scPred* prediction model using scRNA-seq data generated using the SMART-seq2 protocol for 660 dendritic cells. The best fit model from *scPred* used 11 PCs, which collectively explained 5.97% of the variance in the entire training data cohort.

We then applied our model to predict dendritic cells from two independent test data cohorts consisting of scRNA-seq data from a heterogeneous mix of cells from peripheral blood (461 cells) and umbilical cord (420 cells), also generated using the SMART-Seq2 protocol in a different laboratory [33]. Notably, the accuracy for peripheral blood-derived cells was of 98% (Table 2, Figure 3). When we applied the *scPred* model to the cells obtained from an umbilical cord, the overall accuracy was 82%. This lower prediction accuracy possibly reflects a contamination or incorrect original classification of cells obtained from the umbilical cord. To evaluate this, we looked for differentially expressed genes between the 60 cells with a dendritic cell class probability of < 0.9 and the remaining cord cells (Supporting Material Table 3). We identified an upregulation of genes overlapping the T-cell receptor gamma locus: TRGC2, TARP and X06776 (a truncated mRNA from the TRG gamma gene). Additionally, an overrepresentation of myeloid and neutrophil-related biological processes for upregulated genes was identified in these cells (Supporting Material Table 4). All gene ontologies corresponded to myeloid cells, and the presence transcripts from a T-cell specialized locus suggests the presence of T-cells, or alternatively greater heterogeneity in cord-derived cells. Collectively, these results demonstrate that *scPred* is able to accurately predict cell classes using a model trained on data generated in a different laboratory to the test data, without the need to normalize data between sites. This implies that any potential batch effects, or laboratory effects, are not captured in the informative features used to develop the prediction model.

**Table 2:**
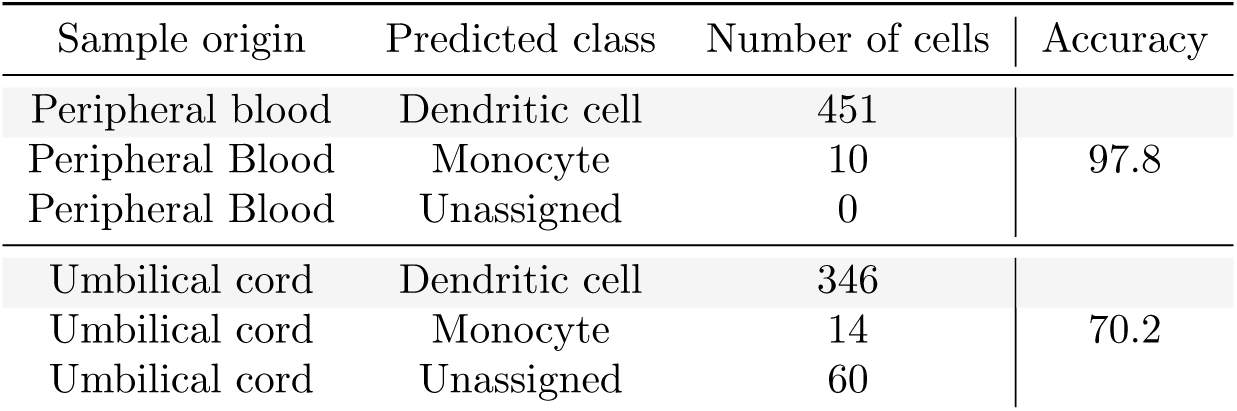
Prediction of dendritic cells from Breton *et al* test data. The first column corresponds to the sample origin of the dendritic cells analyzed by Breton *et al*. The second column shows the class label assigned by *scPred*. The accuracy is reported by sample origin. Importantly, only 10 dendritic cells out of 461 were classified as monocytes. This demonstrates the high accuracy achieved by *scPred* to distinguish dendritic cells from monocytes from peripheral blood. For umbilical cord derived-cells, only 14 out of 420 cells were classified as monocytes and 60 were unassigned as their probability to belong to any of the classes from the training set was low. As discussed in the main text, we argue that these cells correspond to other cell subtypes.

| Sample origin | Predicted class | Number of cells | Accuracy |
| --- | --- | --- | --- |
| Peripheral blood | Dendritic cell | 451 | 97.8 |
| Peripheral Blood | Monocyte | 10 |  |
| Peripheral Blood | Unassigned | 0 |  |
| Umbilical cord | Dendritic cell | 346 | 70.2 |
| Umbilical cord | Monocyte | 14 |  |
| Umbilical cord | Unassigned | 60 |  |

### Accurate classification when cells types are imbalanced

Primary tumors contain cells that both tumor and non-tumor cells of varying types. However, importantly, tumor cells originate from the same cell subtypes of one or more of the original healthy cells in a tissue. Numerous methods exist for classifying (or diagnosing) a whole tissue biopsy as either cancerous or non-cancerous based on DNA genotyping [34], transcriptome profiling [35,36], or histochemistry [37,38]. Most of these methods work well, but are unable to accurately classify heterogeneity at a cellular level, and do not work if the percentage of tumor cells in a biopsy is small. We applied *scPred* to predict epithelial tumor cells from a heterogeneous population of cells from tumor and normal mucosa matched samples from eleven colorectal cancer patients [39]. Of the 275 cells from colorectal cancer samples, the imbalance in the proportions of colorectal cancer epithelial stem/TA-like cells comparted to healthy controls was a 1:5 ratio of normal to tumor cells.

The prediction accuracy was evaluated using a bootstrapping method, training on a randomly sampled 75 percent of the data and predicting on the remaining 25 percent, while correcting for class imbalance [40]. To estimate the variance of prediction accuracy, fifty bootstraps were performed, and the mean across replicates was calculated (Methods). Overall, mean area under the receiver-characteristic function the was 96.4 with a 95% confidence intervals of 95.5 - 97.2 (Figure 5a). Likewise the mean precision-recall curve was 0.992 (95% confidence intervals of 0.989 - 0.995) (Figure 5b). Given the imbalance in the proportions of colorectal cancer, the high area under the precision-recall curve and small confidence intervals indicates that *scPred* is robust to class imbalance in the training data. The high specificity of the model under this scenario implies that a single cell prediction method would be able to accurately diagnosis disease status using scRNA-seq data from a limited number of cells. For example, here the mean sensitivity for tumor cells is 0.761 and the specificity is 0.958. Thus, if in a patient sample 100 cells were single cell sequenced the probability of incorrectly classifying 10 cells as a tumor cells from a healthy individual would be 6.3*x*10^−7^. Conversely, once 10 cells are correctly classified tumor cells in a true tumor biopsy the probability of accurately diagnosing the disease state is approximately 1.

**Figure 5:**
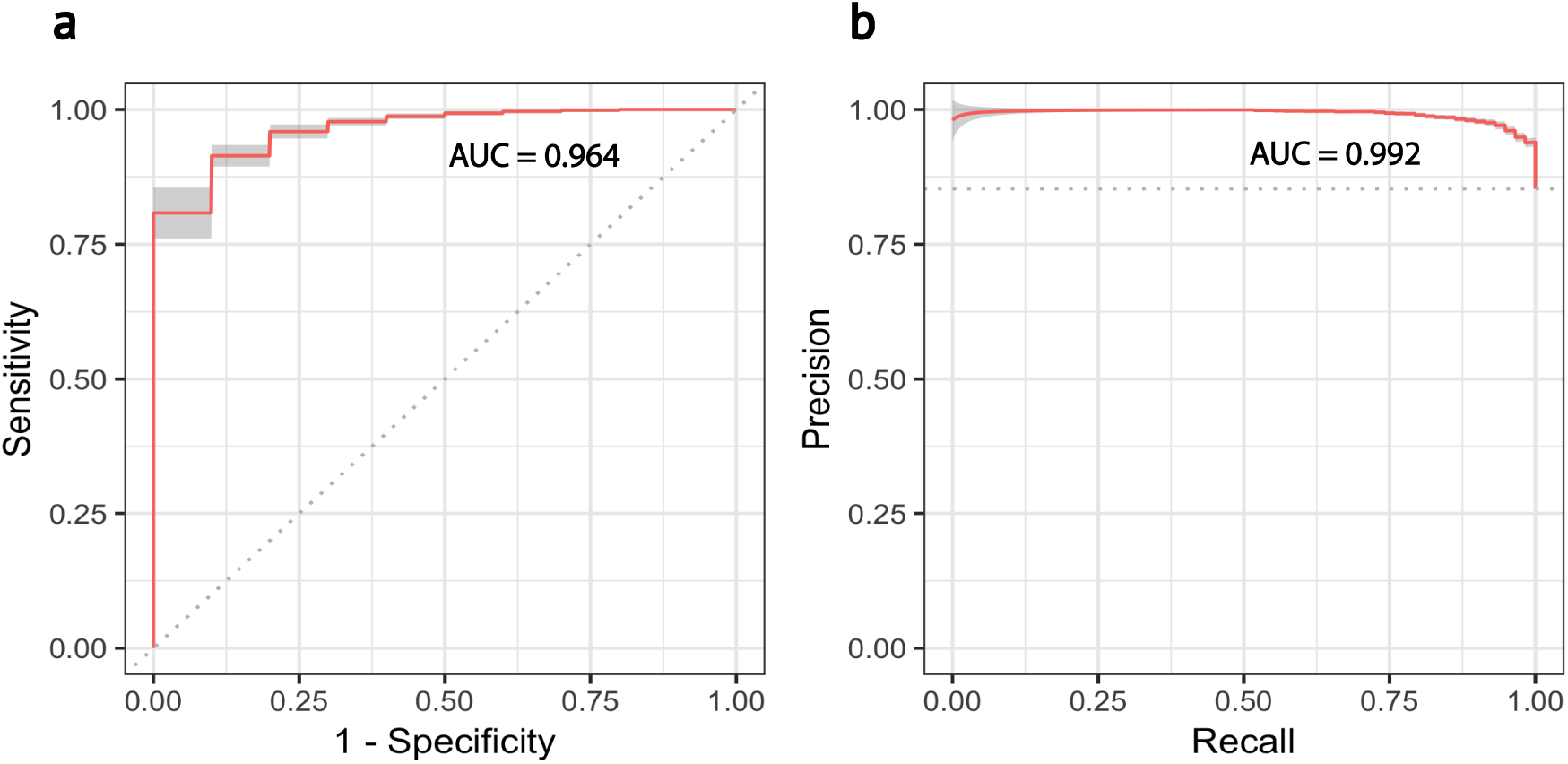
Prediction results of colorectal cancer epithelial stem/TA-like cells. The performance of the prediction was measured using the receiver operating characteristic area under the curve (ROC AUC) and the precision-recall area under the curve (PR AUC). 95% confidence bands are shown in both cases for 50 bootstrap replicates. a) ROC AUC. The area under the curve shows the relationship between the cells incorrectly assigned to that come from tumor samples versus the ones that were correctly assigned by the prediction model as tumor cells using a series of different threshold points. b) PR AUC. The area under the curve measures the relationship between the cells correctly classified as tumor cells versus the fraction of cells correctly assigned as tumor cells from the total number of cells classified as tumor cells. An AUC value of 0.992 shows robustness to class imbalance

### Software

`scPred` is implemented in R as a package based on *S4* objects. The *scPred* class allows the eigendecomposition, feature selection, training and prediction steps in a straightforward and user friendly fashion. `scPred` supports any classification model available from the *caret* package [41]. The default model in *scPred* is the support vector machine with a radial kernel. However, while other models are available we anticipate we would expect their performance to vary based on the distribution of true effects in the fitted PCs (Supporting Material Table 5 and 5). The *scPred* object contains slots to store the eigedecomposition, informative features selected and trained models, meaning models can be applied without re-computing the initial training step. The package also includes functions for exploratory data analysis, feature selection, and graphical interpretation, and can be downloaded from https://github.com/IMB-Computational-Genomics-Lab/scPred/. All analyses were run in a personal computer with 16 GB RAM memory and a 2.9 GHz Intel Core i5 processor.

## Methods

The `scPred` method is split into two major steps. First a prediction model is built using a training dataset of scRNA-seq data. The second step is the application of this prediction model to scRNA-seq data obtained from independent sample, with each cell then assigned a probability of belonging to a given class based on the fit of its scRNA-seq expression levels in the prediction model. Below, we have outlined the methods for each of these steps. We start with a single cell gene expression matrix *C_Train_* (CPM values - Count Per Million Mapped Reads) obtained from different characterization classes: for example, from different cell subtypes, cells obtained from disease verses control samples, or cells defined as different states.

### Training step

The training expression matrix is log2-transformed log_2_(*C_Train_* + 1) to linearize the expression values for each gene and stabilize the variance across a large expression range. Let *G_Train_* be the log_2_-transformed expression matrix *C_Train_* with *n* single cells and *m* genes,

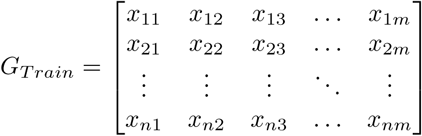

We subsequently center and scale *G_Train_* using the mean and standard deviation of gene expression values of each gene, calculated using the following formulas,

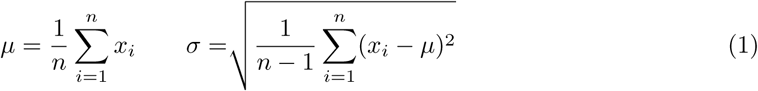

Each mean is subtracted from all *m*th elements of their corresponding *n*th row, and the result is divided by the respective standard deviation as follows:

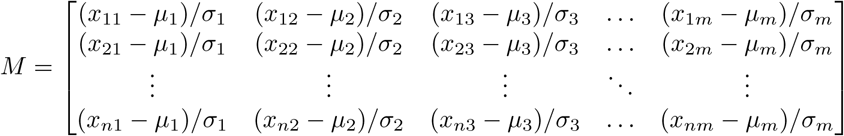

We next calculate orthogonal vectors for the gene expression values using a Singular Value Decomposition (SVD) method. To do so, the matrix *M* needs to be factorized into the product of three matrices as follows:

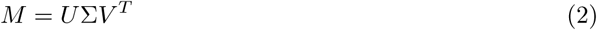

Where *U* and *V* are orthonormal matrices and Σ a diagonal matrix.

First, we compute the product *MM^T^*. To find *U*, we orthogonally diagonalize *MM^T^*:

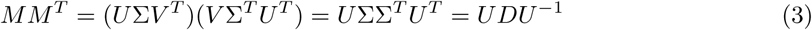

Then, *U* contains the eigenvectors of *MM^T^* (or left singular vectors of *M*) and *D* its eigenvalues.

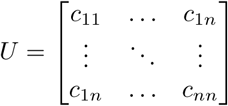

Similarly, to calculate *V*, we compute the product *M^T^ M* and diagonalize *M^T^ M* to calculate its eigenvectors and eigenvalues.

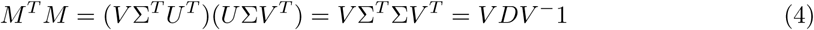

*V* contains the eigenvectors of *M^T^ M* (or right singular vectors of *M*) and *D* its eigenvalues.

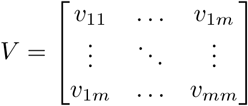

Σ is a diagonal matrix with the squared root eigenvalues of *M^T^ M* (or singular values of *M*) along the diagonal.

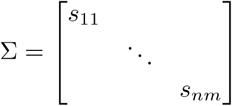

The matrix product *U*Σ gives the principal components (PCs) or “scores”, which are a new set of uncorrelated linear variables that capture the maximum variance from the single cell expression matrix *M*. The individual squared values of the diagonal entries of Σ divided by the sum of all squared values give the variance explained by each principal component. PCs are in descending order according to the variance each of them explains.

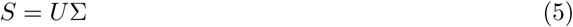

We next identify the PCs whose scores have significant differences between the classification cell classes. We initially create a subspace of *S* (namely *R* with *n* rows and *r* columns (dimensions)), such that each dimension explains at least 0.01% of the variance of the matrix *M*. However, it is important to note that at this stage we do not select features to fit in a prediction model. To identify the informative dimensions, a two-tailed Wilcoxon rank sum test is performed for each PC to assess whether there is a significant difference in the distributions of PC scores for cells in different classes. The resulting p-values are adjusted for multiple testing using a Benjamini-Hochberg false discovery rate correction. Columns from *R* are ranked in ascending order based on their corresponding p-values. This step allows us to identify PCs with the largest difference in their distributions of the scores between the classes, and thus are expected to be the most informative as features used as input predictors in a classification model.

From *R*, we create a subspace *F* with only *f* columns with associated adjusted p-values less than 0.05. The columns of *F* are used as features to train a support vector machine model with a radial kernel [42]. A support vector classifier consists of a subspace (called hyperplane) of dimension *h* − 1 with regard to its ambient high dimensional space *H* with *h* dimensions, which linearly separates the observations (cells) according to the class they belong to (see equations 6 and 7). A margin around the hyperplane is defined in order to minimize the misclassifications. The width of the margin is determined by observations called *support vectors.* Here, we find a hyperplane that separates single cells based on their PC scores into the classification classes. Those cells that define the margin can be thought as *supporting cells* of the hyperplane.

When the observations cannot be separated in the feature space using a linear boundary, a “kernel trick” is used to map observations into a high dimensional space where they can be linearly separated by a hyperplane. Let Φ be a function that maps single cells from a *F* space of *f* dimensions to a higher dimensional space *H*.

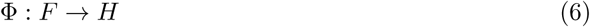

And *k*(*x, x_i_*) be a kernel function that returns the inner product of the images of two cells (based on the values of the *f* principal components in *F*).

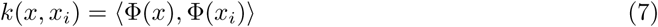

However, instead of computing a feature map Φ for all observations, the following shortcut is possible using a Gaussian radial basis kernel [42]:

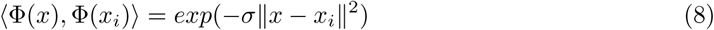

where *σ* is a constant greater than zero estimated via cross-validation. Thus, equation (7) can be rewritten as:

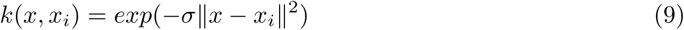

Hence, the coordinates of the cells in *H* are not computed.

Then we can define a function *f*(*x*) that returns a decision value which indicates whether a cell belongs to a class or the other using the kernel function.

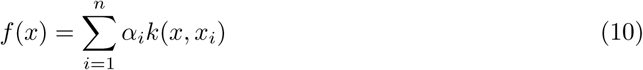

*α_i_* parameters are estimated by solving the following minimization problem:

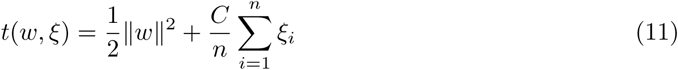

subject to

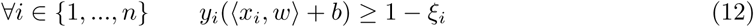

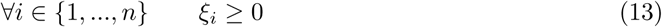

And being the hyperplane defined by the following set:

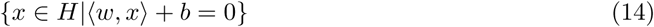

*w* is a weight vector in the feature space *N* perpendicular to the hyperplane which helps to define the margin, *ξ* is a slack variable that allows each cell to be on the wrong side of the hyperplane or the margin in order to deal with outliers, *n* is the number of observations (cells), *y_i_* is a variable that indicates whether the cell *x_i_* belongs to one class (*y* = 1) or the other (*y* = −1) and *C* is a cost parameter that penalizes the sum of *ξ_i_*. As *C* increases, the margin becomes wider and more tolerant to violations by cells. By enlarging the feature space using a polynomial kernel, the cells are linearly separated in *H* [43]. To train the model, we determine the cost *C* and *σ* parameters via cross-validation and select the values that maximize the prediction performance. Finally, class probabilities are calculated using a sigmoid function fitted on the decision values returned by the classifier *f* (*x*) [42].

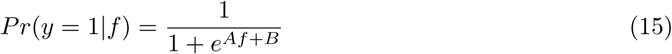

The final trained model consists of a set of parameters that maximizes the margin between the training observations and the hyperplane in order to separate single cells according to their classification class. K-fold cross-validation is performed as described in the `caret` [41] package. If the number of classes is more than two for the response variable, then n binary classification models are trained. For each classification model, we categorized all cells into two classes depending on the class being studied: *positive class* (cell type(s) of interest) and *negative class* (remaining cell types), *one-versus-all* approach.

### Prediction step

Once the model has been trained and evaluated, it can be used to classify single cells from an independent dataset from which the cell classes are unknown. Here, we apply the trained model(s) to classify cells from a testing dataset.

Given a test expression matrix *C_Test_* with *n* single cells as rows and *m* genes as columns, let *G_Test_* be the log_2_-transformed expression matrix *C_Test_*

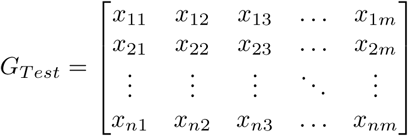

the matrix is centered and scaled using the means and variances calculated from *G_Train_*,

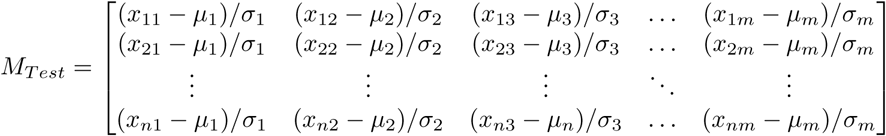

and *M_Test_* is projected onto the training PCA coordinate basis using the rotation matrix *V* after log_2_-transforming and scaling the data according to the training feature space:

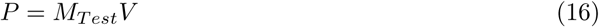

*P* contains the projection of the single cells from test dataset onto the PCs from the training data. Informative PCs listed in the *R* training subspace are extracted from *P* and used as features to predict the classification classes of the cells from the test dataset using the trained support vector machine model (Figure 1). If more than two models were trained, all cells in *P* are classified using the *c* trained models. If the maximum probability obtained across all models is greater than a threshold (0.9 by default), the cell is labeled according to the positive class corresponding to model the highest probability, otherwise the cell is labeled as “Unassigned”.

### Predicting cell type from scRNA-seq data using scPred

Our `scPred` method provides a generalized framework to classify a given cell based on its gene expression values. Importantly, our method is designed to solve the problem of individual gene feature selection and enable subtle effects spread across many genes to be utilized through orthogonal components of variance. In doing so we anticipate an increase in the prediction performance over current gene-centric feature selection, as `scPred` will incorporate the small effects of many genes. To demonstrate both the utility and performance of `scPred` we first validated the performance against an orthogonal molceular asay, and then addressed three distinct biological examples of classification of single cells. Firstly, by predicting specific *α*, *β*, *δ* and *γ* cell subtypes from pancreas islets of Langerhans. Secondly, classifying dendritic cells using a heterogeneous mix of single cells as a reference. And finally, identifying the presence of cancer cells from a heterogeneous composition of cells from whole tissue in both tumors and matched healthy controls.

For all datasets, we removed all cells above or below 3 median absolute deviations (MAD) from the median library size, mitochondrial and ribosomal gene expression. Furthermore, all genes with zero counts across all cells and genes not expressed in at least 1% of the whole population were discarded. Finally, all count matrices were transformed to CPM values and genes being expressed more than five CPM were preserved.

### Gastric cancer tumor verse non-tumor prediction

The collection of this data was conducted in compliance with the Helsinki Declaration. The institutional review board at Stanford University School of Medicine approved the study protocol (19071), and informed consent was obtained. We collected a matched set of samples including a gastric primary cancer, and normal stomach tissue. Tissue biopsies were obtained from surgical resection of a primary gastric adenocarcinoma and matched adjacent normal tissue. Immediately after resection, the tumor sample was stored in RPMI medium on ice for less than one hour. The samples were then macrodissected and dissociated into a cellular suspension by the gentleMACS Octo Dissociator as per manufacturer’s recommendations and the 37C_h_TDK_3 program (Miltenyi Biotec, Bergisch Gladbach, Germany). Single cell RNA-Seq was performed after thawing cryopreserved sample stored in liquid nitrogen in DMSO. Histopathology of this gastric cancer revealed moderate to poorly differentiated features with a 60-70% tumor fraction. Immunohistochemistry demonstrated a loss of MLH1 and PMS2 expression. The loss of these proteins indicated that this tumor had microsatellite instability (MSI) where cancer cells have a hypermutable state because of loss of DNA mismatch repair. The tumor tissue was disaggregated into a single cell suspension and analyzed scRNA-seq.

We used the Chromium Controller instrument (10X Genomics Inc., Pleasanton, CA) and the Single Cell 3’ Reagent kit (v2) to prepare individually barcoded scRNA-Seq libraries following the manufacturer’s standard protocol. Briefly, single cell suspensions were loaded on a Chromium and were partitioned in droplets. Reverse transcription is performed, followed by droplet breaking, and cDNA amplification. Each cDNA molecule thus contained the read 1 sequencing primer, a 16bp cell-identifying barcode, and a 10bp UMI sequence. We performed enzymatic fragmentation, end-repair, and a-tailing followed by ligation of a single-end adapter containing the read 2 priming site. Finally, sequencing libraries were quantified by qPCR before sequencing using 26x98 paired-end reads. The Cellranger software suite was used to process scRNA-seq data, sample demultiplexing, barcode processing, and single cell 3’ gene counting. Cellranger provided a gene-by-cell matrix, which contains the read count distribution of each gene for each cell.

The gene expression matrix was split in halves according to the disease status of each cell (tumor or normal) to create a training and testing dataset. We selected only class-informative PCs explaining at least 0.01% of the variance and using an adjusted alpha threshold of 0.05. 10-fold cross-validations were performed to train a support vector machine model with a radial kernel. This procedure was repeated ten times as bootstrap replicates. Finally, we obtained the sensitivity and specificity to measure the performance of `scPred`.

### Prediction of Islets of Langerhans subtypes

We considered three independent datasets to train a prediction model to classify *α* (alpha), *β* (beta), *δ* (delta) and *γ* (gamma) cell subtypes: Muraro *et al.* [24] consisting in 1,522 cells from a CEL-Seq2 protocol. Segerstolpe *et al.* [3] consisting of scRNA-seq data from 1,321 cells generated using the Smart-Seq2 protocol and, 1,349 cells from Xin *et al* [25]) whose gene expression levels were assayed using the SMARTer protocol (Supporting Material Table 1). We integrated the three datasets using the intersection of genes between them and obtained a single aggregated matrix.

We applied the Seurat alignment approach [29] to account for technical differences across the different datasets used for the training dataset. First, we determined the most variable genes (528) in at least two of the three datasets to compute the loadings and the first 30 PCs using the implicitly restarted Lanczos bidiagonalization algorithm [44]. Then, we used the loadings from the training eigendecomposition to project the testing dataset (Baron *et al*.) and obtained the cell embeddings. After the alignment, no batch effect was observed (see Figure 3).

Then, we trained a prediction model considering only class-informative PCs using a multiple testing corrected *alpha* level of 0.05 using (see `scPred` methods section) using the scores from the aligned training eigenspace only. 10-fold cross-validations were performed to train a support vector machine model with a radial kernel. To assess the performance of our prediction model, we predicted the specific cell types of 7,932 cells using their scRNA-seq data generated using the inDrop protocol [30].

Muraro, Xin, and Baron datasets were obtained from Gene Expression Omnibus (GEO) under the accession numbers GSE85241, GSE81608, and GSE84133 respectively. Segerstolpe dataset was obtained from the Array Express under accession number E-MTAB-5060.

### Prediction of dendritic cells

Training data of dendritic cells and monocytes scRNA-seq data (processed using a Smart-Seq2 protocol) [1]. After quality control, 660 dendritic cells and 335 monocytes were used to train a prediction model applying a 0.01% variance-explained filter and a corrected alpha level of 0.05 to select the informative PCs. 10-fold cross-validations were performed to train a support vector machine model with a radial kernel. We tested the `scPred` prediction model against dendritic cells from an independent study [33], whose scRNA-seq data had been generated using the SMART-Seq 2 protocol. After quality control, 150 primary human conventional dendritic cells (cDCs), 420 cord blood pre-cDCs and 311 blood pre-cDCs were kept.

The final training model consisted of only 11 discriminant PCs explaining 5.97% of the variance from the Villani *et al.* dataset. The training error of the model was 0.018 and 232 cells were used as support vectors. Differential expression analysis between the unassigned cells and remaining cells from cord blood was performed using edgeR [45]. Genes with a log fold-change greater or lower than 2 and an adjusted p-value less than 0.05 were considered as differentially expressed. Gene ontology analysis was performed using http://pantherdb.org/.

Training data (Villani *et al*.) was obtained from (https://portals.broadinstitute.org/single_cell) and test dataset (Breton *et al*.) from the Gene Expression Omnibus (accession number GSE89232).

### Prediction of colorectal cancer cells

We obtained the human colorectal cancer dataset under the GEO accession number GSE81861. We analyzed only the stemTA cell subtype as they are the most abundant epithelial subpopulation. After quality control, 275 cells stemTA cells and 21,933 genes were kept.

The gene expression matrix was split according to the sample origin of each cell (tumor or normal) to create a training and testing dataset such that the former partition contained 75% of cells and the latter 25%. To create the partitions, we used the SMOTE algorithm [40] to account for class imbalance. We selected only class-informative PCs explaining at least 0.01% of the variance and using an adjusted alpha threshold of 0.05. 10-fold cross-validations were performed to train a support vector machine model with a radial kernel. Finally, we obtained the areas under the ROC and precision-recall curves to measure the performance of `scPred`. Ten bootstrap replicates were performed.

## Discussion

Single-cell RNA sequencing has provided the ability to analyze the transcriptomic profile of individual cells, leading to the identification of novel cell types and the characterization of heterogeneous cell populations. Here we introduced `scPred`, a novel method to classify single cells based on singular value decomposition and a support vector machine model. `scPred` takes advantage of the informative signals spread across orthonormal linear combinations of the gene expression values and minimizes the incorporation of noise to the prediction model by excluding principal components with low contribution to the variance explained. `scPred` uses support vector machines as a default machine learning approach as it is suitable for large datasets and accounts for various sources of data [46].

Collectively, our results show that *scPred* is able to accurately classify individual cells from an independent sample to those used to train the prediction model. However, the ability to do so even when using a training cohort of cells whose scRNA-seq data is assayed from different platforms, has important implications for a practical implementation of *scPred.* The ability to build a single cell training cohort using data generated from multiple platforms means that composite reference datasets can be generated, which will increase the predictive accuracy of *scPred* through more accurate estimate of the model effects.

One of the advantages of `scPred` is that by reducing the dimensions of the gene expression matrix via singular value decomposition we also decrease the number of features to be fit, reducing both the computational requirements for prediction and the prediction model parameter space. While we have used a support vector machine method, the `scPred` software can be easily adjusted to use other classification algorithms [41], allowing a user to choose the models that suit the effect distributions of their data best.

---

**Table S1.**
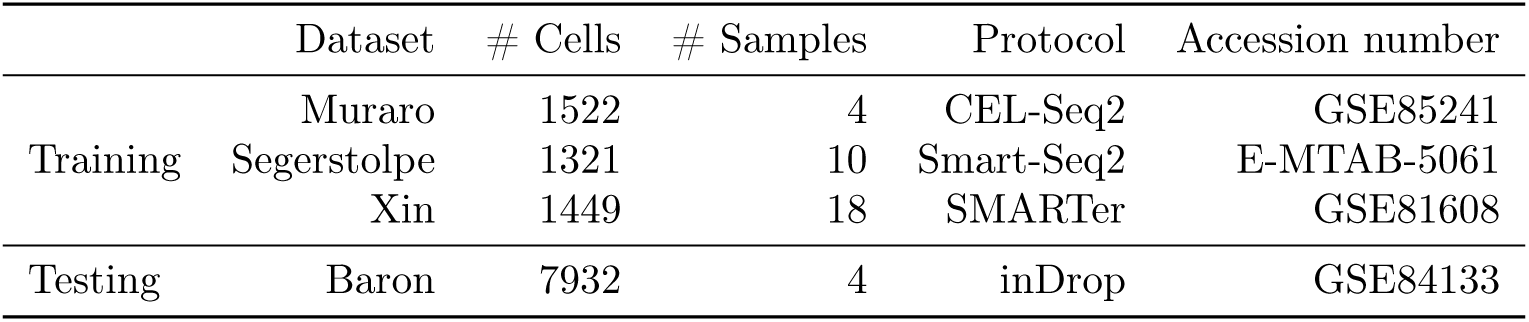
Summary of pancreas datasets. Training dataset consisted of 4,292 cells and 32 human samples in total. All datasets were generated using different protocols. All four samples from the Muraro dataset derive from healthy individuals, as well as 6 samples from the Segerstolpe dataset and 12 from Xin. All the remaining 10 samples from the training reference and 4 from the testing phase come from diabetic individuals. The incorporation of 32 individuals -both healthy and diabetic- to train the prediction model captured a broad biological variability to assess the cell identity of pancreatic cells in other datasets.

|  | Dataset | # Cells | # Samples | Protocol | Accession number |
| --- | --- | --- | --- | --- | --- |
| Training | Muraro | 1522 | 4 | CEL-Seq2 | GSE85241 |
|  | Segerstolpe | 1321 | 10 | Smart-Seq2 | E-MTAB-5061 |
|  | Xin | 1449 | 18 | SMARTer | GSE81608 |
| Testing | Baron | 7932 | 4 | inDrop | GSE84133 |

**Table S2:** Prediction results of pancreatic cells from Baron dataset. The first column shows the predicted classes by `scPred` and the remaining columns the true classes. Values along the diagonal corresponds to the number of cells that were correctly classified. The “Unassigned” label is used by `scPred` when a cell cannot be classified with confidence as Alpha, Beta, Delta or Gamma. The “Other” column comprises other cell types except cells from the islets of Langerhans.

| Prediction | Alpha | Beta | Delta | Gamma | Other |
| --- | --- | --- | --- | --- | --- |
| Alpha | 2264 | 2 | 0 | 1 | 32 |
| Beta | 1 | 2359 | 0 | 0 | 60 |
| Delta | 1 | 13 | 579 | 0 | 21 |
| Gamma | 3 | 2 | 0 | 252 | 5 |
| Unassigned | 33 | 78 | 14 | 1 | 2208 |

**Table S5:**
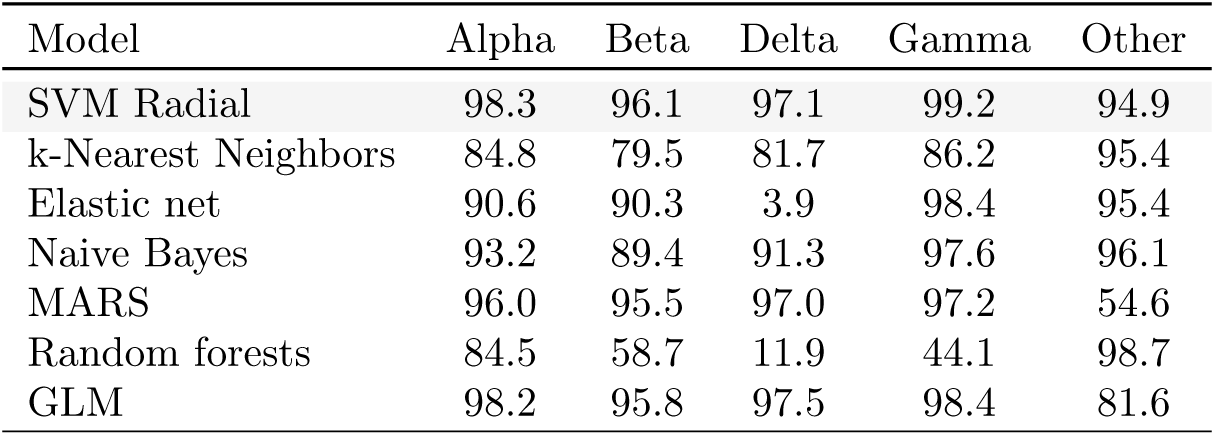
Prediction performance of pancreatic cells from *Baron et al.* dataset using different prediction models described in table S1. Using a threshold of 0.9 to define class identity for each cell, the support vector machine model with a radial kernel performed better compared to other models as the prediction results show high specificity (for other cells) and high sensitivity (for cell types from the islets of Langerhans).

| Model | Alpha | Beta | Delta | Gamma | Other |
| --- | --- | --- | --- | --- | --- |
| SVM Radial | 98.3 | 96.1 | 97.1 | 99.2 | 94.9 |
| k-Nearest Neighbors | 84.8 | 79.5 | 81.7 | 86.2 | 95.4 |
| Elastic net | 90.6 | 90.3 | 3.9 | 98.4 | 95.4 |
| Naive Bayes | 93.2 | 89.4 | 91.3 | 97.6 | 96.1 |
| MARS | 96.0 | 95.5 | 97.0 | 97.2 | 54.6 |
| Random forests | 84.5 | 58.7 | 11.9 | 44.1 | 98.7 |
| GLM | 98.2 | 95.8 | 97.5 | 98.4 | 81.6 |

**Table S6:**
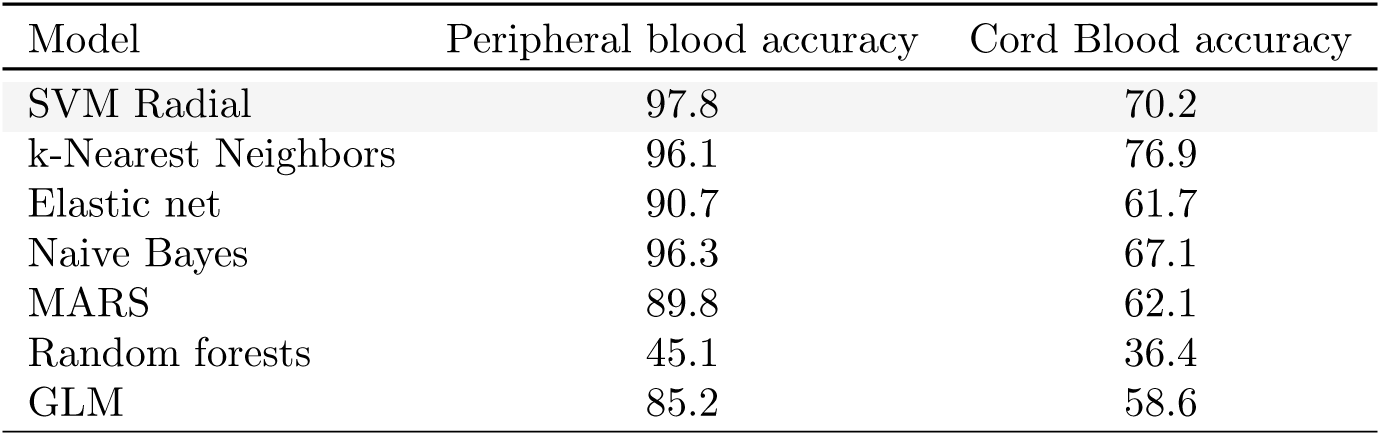
Prediction of dendritic cells from *Breton et al.* dataset using different prediction models. Except from random forests, all models showed a high accuracy for dendritic cells from peripheral blood. For cord blood-derived cells, wide differences are observable across models due to the presence of other subpopulations. The support vector machine model showed the best accuracy for peripheral blood-derived dendritic cells. Accuracy is defined as the fraction of real dendritic cells correctly predicted by `scPred`.

**Table S3:** Differentially expressed genes between unassigned cells by scPred and remaining cord blood-derived cells. Top upregulated genes include T-cell receptor gamma locus and myeloid-related genes. See spreadsheet excel file.

**Table S4:** Gene ontology overrepresentation results of overexpressed genes from unassigned cells. A Fisher’s exact with FDR multiple test correction. Biological processes involving myeloid and neuthophiles were overrepresented. X06776, XIST, BC039116, M64936, TARP, FCGR1C, and ECRP gene identifiers did not map the query database. See spreadsheet excel file.

**Fig S1.**
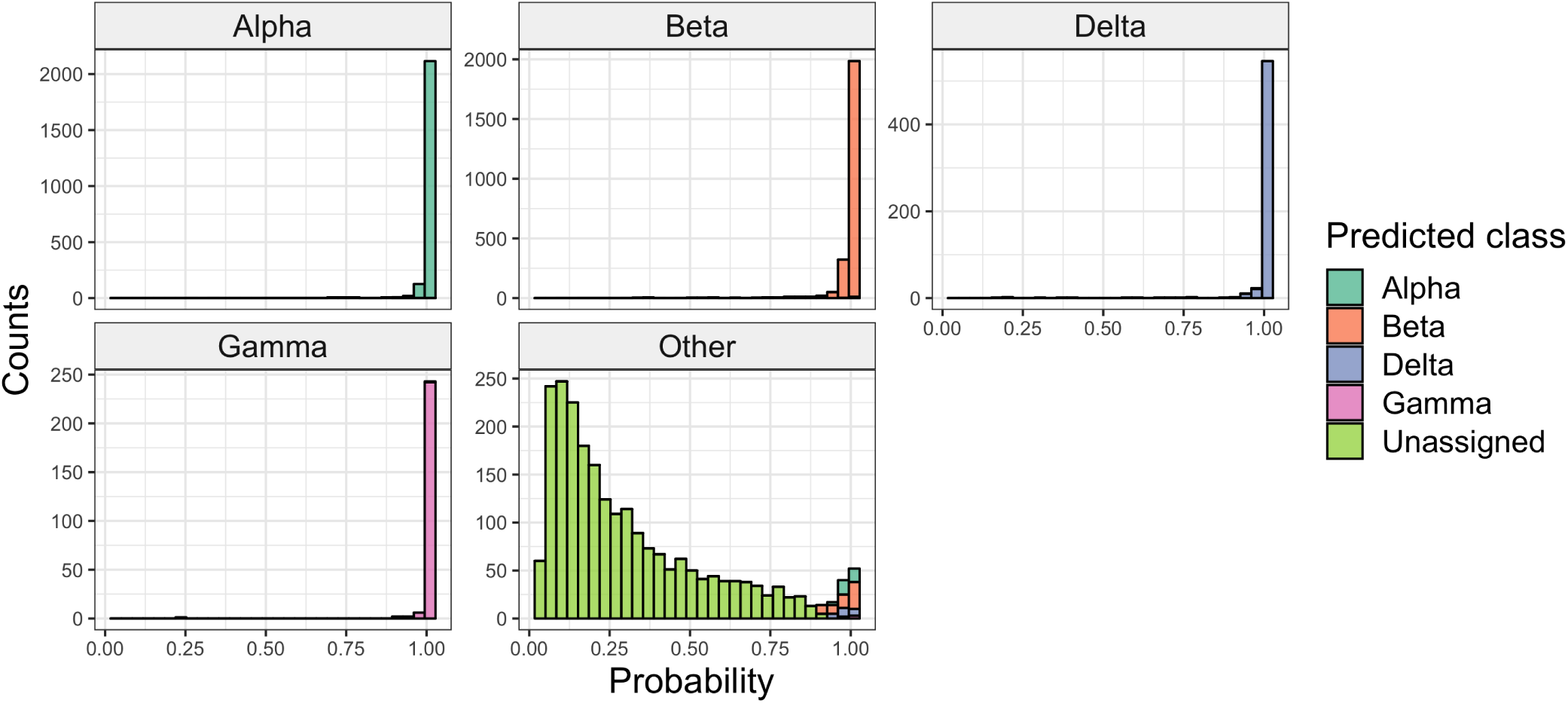
Distribution of conditional class probabilities for single cells from the Baron test dataset across all four models. Each panel corresponds to the true cell type classes and each color to the predicted class by `scPred`. The right-skewed distributions for *α*, *β*, *δ*, and *γ* cells indicate a high confidence prediction for most cells from the Islets of Langerhans. The left-skewed distributions for “Other” cells suggests that most of these cells are not likely to belong to any of the cell types of interest.

